# Risk perceptions and preventive practices of COVID-19 among healthcare professionals in public hospitals in Ethiopia

**DOI:** 10.1101/2020.11.04.367896

**Authors:** Wakgari Deressa, Alemayehu Worku, Workeabeba Abebe, Muluken Gizaw, Wondwosson Amogne

## Abstract

Healthcare professionals are at higher risk of contracting the novel coronavirus due to their work exposure in the healthcare settings. Practicing appropriate preventive measures to control COVID-19 infection is one of the most important interventions that healthcare workers are expected to use. The aim of this study was to assess the level of risk perception and practices of preventive measures of COVID-19 among health workers in Addis Ababa, Ethiopia. A hospital-based cross-sectional study was conducted from 9^th^ to 26^th^ June 2020 among healthcare professionals working at six public hospitals in Addis Ababa. Data were collected using a self-administered structured questionnaire. Frequency, percentage, and mean were used to summarize the data. A binary logistic regression analyses were performed to identify factors associated with risk perception about COVID-19. A total of 1,134 participants were surveyed. Wearing facemask (93%), hand washing for at least 20 seconds (93%), covering mouth and nose while coughing or sneezing (91%), and avoiding touching eyes, nose, and mouth (91%) were the commonly self-reported preventive practices. About 88% perceived that they were worried about the risk of becoming infected with coronavirus, and majority (91%) worried about the risk of infection to their family. The mean score of overall fear and worry of COVID-19 was 2.37 on a scale of 1 to 3. Respondents who ever provided clinical care to COVID-19 patients were more likely to report fear and worry (adjusted OR=1.34, 95% CI:1.02-1.91), however those who ever participated in Ebola or SARS outbreaks were less likely to report fear and worry due to COVID-19 crisis (adjusted OR=0.66, 95% CI:0.48-0.90). This study has revealed widespread practices of preventive measures and the highest perceived risk of COVID-19 among healthcare workers. Therefore, an effective risk communication intervention should be implemented to ensure the maintenance of appropriate practices during the current COVID-19 pandemic.

## Introduction

The novel coronavirus disease 2019 (COVID-19) that was declared as a pandemic by the World Health Organization (WHO) on the 11^th^ of March 2020 [1] has affected over 37 million people and has caused more than one million deaths globally as of 12^th^ October 2020 [2]. The new severe acute respiratory syndrome coronavirus 2 (SARS-CoV-2) has now spread to 213 countries and territories around the world. Up to 20^th^ September 2020, Ethiopia reported a total of 68,820 confirmed coronavirus disease 2019 (COVID-19) cases and 28,314 recoveries from over 1,202,818 total tests, among whom 1,096 have died [3]. Over 1,311 health workers have contracted coronavirus in Ethiopia as of 17^th^ September 2020.

Healthcare providers who are in the healthcare settings to care for the COVID-19 patients are highly vulnerable to SARS-COV-2 infection [4]. Most healthcare workers are working in isolation units, critical care units, intensive care units (ICUs), emergency units, working in frontline positions, and having contact with suspected and confirmed COVID-19 cases. During the early stage of COVID-19 pandemic in the USA, the prevalence of SARS-CoV-2 infection among healthcare workers was 7.3% and particularly, infections were most common among nurses [5]. In the south of the Netherlands, 96 (5%) of 1796 health care workers screened in three hospitals were tested positive for SARS-CoV-2 just 10 days after the first reported COVID-19 case in the country [6]. More than 278 physicians from almost all medical specialties have died due to COVID-19 as of 15 April 2020 with the majority (44%) from Italy mainly because of lack understanding of the virus and its preventive measures [7]. Studies in China reported 3,387 COVID-19 cases among HCWs (4.4% of all cases), with 23 attributable deaths [8]. In some countries at the peak of their infection, such as Spain, they have reported that 13% to 14% of the country’s cases were in healthcare workers [9]. Overall, as much as 10% of healthcare workers are infected with SARS-CoV-2 in some countries [4] and the WHO has developed infection prevention and control guidance to be implemented at the national and healthcare facility level in order to reduce coronavirus infection among healthcare workers [10].

Studies have identified major sources of worry and anxiety among healthcare professionals due to lack of appropriate PPE; being exposed to COVID-19 at work and taking the infection home to their family; not having rapid access to testing if they develop COVID-19 symptoms and concomitant fear of propagating infection at work; uncertainty that their organization will support/take care of their personal and family needs if they develop infection; access to childcare during increased work hours and school closures; and support for other personal and family needs as work hours and demands increase [11]. A recent qualitative study from China reported the challenges facing frontline healthcare workers during the COVID-19 outbreak, including a high risk of infection, insufficient PPE, heavy workloads and manpower shortages, confusion, discrimination, isolation, separation from their families, and burnout [12]. Under these stressful conditions, healthcare professionals have been challenged to effectively engaged in the fight COVID-19.

A good level of understanding the risk perception and preventive practices of healthcare professionals is essential to protect the health workers and prevent the COVID-19 pandemic through effective risk communication. Studies conducted during the early stages of a pandemic have suggested that perceived personal risk of infection and the health effects are linked to engagement in protective behaviors [13]. Since the occurrence of the epidemic in Ethiopia, the MoH, in collaboration with its partners, conducted different trainings on preventive measures for healthcare professionals at several hospitals and health centers, with supplies of PPE materials. However, so far, no study has been undertaken in Ethiopia on risk perception and preventive practices of healthcare professionals during the current COVID-19 pandemic. In addition, levels of confidence and feelings of healthcare workers about COVID-19 are unknown. It was therefore necessary to carry out this study to investigate the level of risk perception and preventive practice of healthcare professionals towards the COVID-19.

## Methods

### Study setting and design

This hospital-based cross-sectional study was conducted from 9^th^ to 26^th^ June 2020 at six public hospitals in Addis Ababa city administration, three months after the first confirmed COVID-19 case in Ethiopia in March 2020. Addis Ababa city is the most populated urban city in the country, and had a population of about 3.6 million in 2019 [14]. The city also had better health infrastructure and the highest number of qualified medical personnel compared with any city or region in the country. There were 12 hospitals and close to 100 health centers belonging to the public center, and about 25 private hospitals in Addis Ababa city. There were also over 17,000 healthcare professionals in the city, including 2,441 (14%) physicians and 8,172 (47%) nurses by the end of July 2019 (MOH 2011 EC Health Indicators). The hospitals selected for the current study provide outpatient and inpatient services for the city residents and patients coming from different parts of the country.

### Study population and sampling

The study was conducted among all healthcare professionals working in the different clinical departments or units of six public hospitals in Addis Ababa, mainly Gyn&Ob, Surgery, Pediatrics, Internal Medicine, OPD, emergencies, intensive care, operation room/ward, screening/triage, laboratory and anesthesia. The selected hospitals included: Tikur Anbessa Specialized Hospital (TASH), Zewditu Memorial Hospital (ZMH), Ghandi Memorial Hospital (GMH), Menelik II Hospital, Yekatit 12 Hospital Medical College (Y12HMC) and St. Paul Hospital Millennium Medical College (SPHMMC). The study population included intern doctors, resident doctors, general practitioners, medical specialists and sub-specialists, health officers, anesthetists, nurses, midwives, pharmacists, laboratory technologists, physiotherapists, X-ray and laboratory technicians, all of whom may expect to encounter suspected or confirmed COVID-19 patients.

A multi-stage sampling, using a mix of purposive and non-random sampling, was applied to select the study participants. In the first stage, the six hospitals were purposively selected from 12 hospitals in the city. In the second stage, clinical departments or units were selected, and in the third stage, study participants were selected proportionally to the estimated number of healthcare professionals working in different departments and units of the hospital. All eligible participants in each department/unit who consented to participate were recruited into the study. Since COVID-19 is a new disease, we assumed that at least 50% of study participants had higher risk perception regarding COVID-19, and the estimated sample size was calculated with 95% confidence limit, with 4% precision and a design effect equal to 1.5 using 20 % non-response rate. Accordingly, the minimum total sample size targeted for this survey was 1,080 respondents. A total of 1,200 participants were targeted for the study.

### Data Collection

A structured paper-based self-administered questionnaire was used to collect the data. The questionnaire is composed of parts on the demographic (gender, age) and occupational characteristics of the respondents (hospital, department/unit, professional category, and work experience), as well as their preparedness to combat COVID-19, potential risk of becoming infected with the virus, worries about the potential risk to their family and loved ones, feelings and fears about COVID-19. Questions related to measures taken to prevent infection from the virus included hand washing for at least 20 seconds, use of disinfectants, wearing facemask, physical distancing, covering mouth and nose while coughing and sneezing and other preventive measures. The questionnaire was developed in English by the authors of the study based on the previously conducted studies and visiting the WHO websites for frequently asked questions on risk perception of healthcare professionals. Most of the questions were designed as ‘yes/no’, ‘agree/disagree’, and ‘worried/not worried’ using different rating scales.

A total of 12 experienced data collectors with health backgrounds were involved in the data collection of this survey. A guideline was developed by the research team to guide the data collectors and supervisors for data collection, quality assurance of data and ethical conduct. Training and orientation on the survey tool and methodology including how to administer the SAQ were conducted for the data collectors using webinar on 2^nd^ June 2020. After explaining the purpose of the study and obtaining written or oral informed consent, study participants were given a paper-based questionnaire at their workplace and they filled out their own questionnaires. The purpose of the study was clearly stated in the questionnaire and the participants were asked to complete the questionnaire with honest answers after giving their consents. The study participants were encouraged to fill out the questionnaire whilst the data collectors were still in the hospital during the data collection period. A collection center was also prepared in the Hospital Director’s office to also gather the questionnaires from the healthcare workers that were unable to directly deliver the completed questionnaires to the data collectors. The data collection took place simultaneously in the six hospitals. The questionnaires were checked for completeness and consistency upon collection. All responses were anonymous.

Risk perception among the healthcare professionals in this study was measured using questions on perceived fears and worries, vulnerability and feelings, and behavioral responses regarding COVID-19 [15–16]. Preventive practices of COVID-19 in this study include hygiene behaviors (such as hand washing; covering mouth and nose with a hand or tissue while coughing or sneezing; avoiding touching eyes, nose and mouth with unwashed hands; using hand sanitizer; disinfecting surfaces); mask wearing, physical distancing and avoiding crowds and public places [17].

### Data analysis

Data were entered into the Census Surveys Professional (CSPro) Version 7.2 statistical software package and subsequently exported to SPSS version 23.0 (SPSS Inc., IBM, USA) for cleaning and data analysis. Descriptive analysis was applied to calculate the frequencies, proportions and mean scores, and the results were presented as a proportion for the categorical variables, and as a mean ± standard deviation for the quantitative variables. A Chi-square was used to establish significance and relationship between variables. The study participants were asked 12 questions related to their fears and worries (risk perception) about COVID-19, such as losing someone they love due to the disease, health system overcrowding, mental and physical health, etc., on a 3-point scale, where 1=don’t worry at all, 2=worry somehow and 3=worry a lot. A sum of scores (ranged 12-36) was made and the level was classified into two groups using the Visual Binning in SPSS (low fear/worry ≤29 and high fear/worry >29 score). Univariate odds ratios (crude OR) and multivariate odds ratio (adjusted OR) were derived by using univariate and multivariate logistic regression models, respectively, to identify the main factors associated with healthcare workers high risk perception. Statistical significance was considered for *P*<0.05. The internal consistency (reliability) of the questions was tested by applying Cronbach’s alpha and the Cronbach’s alpha coefficient of the reliability of scale was estimated at 0.91, which is highly acceptable.

### Ethical considerations

The study protocol was reviewed and approved by the Institutional Review Board of the College of Health Sciences at Addis Ababa University (AAU). Permission to undertake this study was obtained from every relevant authority at all levels. Official letters from AAU were written to each hospital to cooperate and participate in the survey. The purpose and significance of the study was introduced to the study participants, and all participants provided written or oral consent before participating in the study. Anonymity and data confidentiality were ensured, and no identifiable data from participants were collected. All study respondents were asked to only fill the questionnaire once to avoid duplication of data and that their participation in the study was entirely on voluntary basis. All personnel involved in the survey received orientation on COVID-19 infection prevention and control measures.

## Results

### Characteristics of study participants

A total of 1,134 (92%) healthcare professionals consented and completed the questionnaires, out of 1,228 possible participants from six public hospitals in Addis Ababa. Among 1,134 healthcare personnel, nearly 40% of them were nurses, followed by physicians (22.4%) and interns (10.8%). Table 1 summarizes the demographic and occupational characteristics of the study participants and their professional affiliation. Among 1,102 respondents reporting gender, 45.9% were males, with females making 51.3% of all respondents. Among 982 participants with available data on age, the mean (±SD) age was 30.3±6.4 years and ranged from 22 to 70 years old, with the majority within the age group of 20-29 years (57.9%) (31.0±5.6 years for physicians, 25.6±3.3 years for interns and 30.7±6.5 years for nurses). Among 252 physicians participated in the study, general practitioners and resident doctors accounted for 44.8% and 42.9%, respectively, while medical specialists and sub-specialists accounted for the remaining 12.3%. About 17% of the respondents represented other professional categories such as anesthetist, pharmacist, health officer, radiographer and laboratory technologist. Majority (17.2%) of the respondents worked in Gny&Ob department, while 13.8% were in surgical department, 13.3% in pediatrics, 13.0% in medical and 10.5% in OPD departments. Most respondents worked as staff for less than 10 years in the hospital (73.2%), and nearly 10% worked for 10 or more years.

**Table 1.** Characteristics of study participants by professional category (n=1134)

| Characteristics | Professional category, n (%) |  |  |  |  | Total, n (%) |
| --- | --- | --- | --- | --- | --- | --- |
|  | *Physician* | *Intern* | *Nurse* | *Midwife* | *Other\** |  |
| Gender (n=1134) |  |  |  |  |  |  |
| Male | 157 (62.3) | 58 (47.2) | 175 (38.6) | 44 (37.6) | 103 (54.5) | 537 (47.4) |
| Female | 95 (37.7) | 65 (52.8) | 278 (61.4) | 73 (62.4) | 86 (45.6) | 597 (52.6) |
| Age group (years) (n=982) |  |  |  |  |  |  |
| 20-29 | 101 (45.9) | 99 (91.7) | 220 (57.0) | 80 (79.2) | 69 (41.3) | 569 (57.9) |
| 30-39 | 106 (48.2) | 8 (7.4) | 119 (30.8) | 14 (13.9) | 70 (41.9) | 317 (32.3) |
| ≥40 | 13 (5.9) | 1 (0.9) | 47 (12.2) | 7 (6.9) | 28 (16.8) | 96 (9.8) |
| Mean (±SD) | 31.0 (±5.6) | 25.6 (±3.3) | 30.7 (±6.5) | 28.3 (±5.7) | 32.6 (±7.5) | 30.3 (±6.4) |
| Median (Range) | 30.0 (22-70) | 25.6 (22-45) | 30.7 (22.57) | 28.3 (22-52) | 32.3 (23-60) | 30.3 (22-70) |
| Department/Unit (n=1134) |  |  |  |  |  |  |
| Gyn&Ob | 27 (10.7) | 31 (25.2) | 36 (7.9) | 97 (82.9) | 4 (2.1) | 195 (17.2) |
| Surgical | 43 (17.1) | 31 (25.2) | 65 (14.3) | 2 (1.7) | 16 (8.5) | 157 (13.8) |
| Pediatrics | 39 (15.5) | 35 (28.5) | 71 (15.7) | 2 (1.7) | 4 (2.1) | 151 (13.3) |
| Medical | 62 (24.6) | 17 (13.8) | 62 (13.7) | 0.0 | 6 (3.2) | 147 (13.0) |
| OPD/Screening/Triage | 16 (6.3) | 2 (1.6) | 83 (18.3) | 6 (5.1) | 37 (19.6) | 144 (12.7) |
| Emergency | 28 (11.1) | 4 (3.3) | 34 (7.5) | 10 (8.5) | 19 (10.1) | 95 (8.4) |
| Anesthesia/OR/IC | 12 (4.8) | 1 (0.8) | 66 (14.6) | 0.0 | 14 (7.4) | 93 (8.2) |
| Other*** | 25 (9.9) | 2 (1.6) | 36 (7.9) | 0.0 | 89 (47.1) | 152 (13.4) |
| Hospital (n=1134)*** |  |  |  |  |  |  |
| TASH | 79 (31.3) | 17 (13.8) | 128 (28.3) | 19 (16.2) | 40 (21.2) | 283 (25.0) |
| ZMH | 39 (15.5) | 36 (29.3) | 54 (11.9) | 15 (12.8) | 33 (17.5) | 177 (15.6) |
| GMH | 17 (6.7) | 7 (5.7) | 51 (11.3) | 21 (17.9) | 19 (10.1) | 115 (10.1) |
| Y12HMC | 35 (13.9) | 12 (9.8) | 48 (10.6) | 15 (12.8) | 42 (22.2) | 152 (13.4) |
| MH | 39 (15.5) | 29 (23.6) | 68 (15.0) | 20 (17.1) | 18 (9.5) | 174 (15.3) |
| SPHMMC | 43 (17.1) | 22 (17.9) | 104 (23.0) | 27 (23.1) | 37 (19.6) | 233 (20.5) |
| Work experience (n=938) |  |  |  |  |  |  |
| <5 | 167 (79.5) | 84 (90.3) | 168 (44.0) | 65 (67.0) | 68 (43.6) | 552 (58.8) |
| 5-9 | 33 (15.7) | 7 (7.5) | 160 (41.9) | 25 (25.8) | 53 (34.0) | 278 (29.6) |
| 10-14 | 5 (2.4) | 2 (2.2) | 29 (7.6) | 4 (4.1) | 21 (13.5) | 61 (6.5) |
| 15-34 | 15 (2.4) | 0.0 | 25 (6.5) | 3 (3.1) | 14 (9.0) | 47 (5.0) |
| Total, n (%) | 252 (22.2) | 123 (10.8) | 453 (39.3) | 117 (10.3) | 189 (16.7) | 1134 (100) |

\*Other: Includes anesthetist, pharmacist, health officer, lab technologist and radiographer.
\*\*Other: Includes Isolation room/ward, Pharmacy, Oncology, etc.
\*\*\*TASH: Tikur Anbessa Specialized Hospital; ZMH: Zewditu Memorial Hospital; GMH:Ghandi Memorial Hospital; Y12HMC: Yekatit 12 Hospital Medical College; MH: Menelik II Hospital; SPHMMC: St. Paul Hospital Millennium Medical College.

### COVID-19 preventive practices

The self-reported prevalence of different preventive measures practiced by healthcare professionals to prevent themselves from coronavirus infection is shown in Table 2. The overall highest practice showed among healthcare participants were wearing facemask (93%), hand washing for at least 20 seconds (92.7%), covering mouth and nose when coughing or sneezing (90.9%), and avoiding touching eyes, nose, and mouth with unwashed hands (90.5%). These measures were commonly reported (>90%) for physicians, intern doctors, nurses and other healthcare professionals except the midwives who reported <90%. A lower percentage of self-reported practices were observed in physical distancing (84.3%), the use of disinfecting surfaces (76.1%), and staying home when feeling cold or sick (64.6%), with similar pattern across the different categories of healthcare workers.

**Table 2.** Self-reported prevalence of preventive measures practiced by healthcare professionals to prevent coronavirus infection by professional category (n=1134)

| Variable | Professional category, % |  |  |  |  | Total, % |
| --- | --- | --- | --- | --- | --- | --- |
|  | *Physician* | *Intern* | *Nurse* | *Midwife* | *Other\** |  |
| Wearing face mask | 95.6 | 95.9 | 90.9 | 89.7 | 94.7 | 93.0 |
| Hand washing for at least 20 seconds | 95.2 | 95.1 | 90.9 | 88.9 | 94.2 | 92.7 |
| Covering your mouth and nose when you cough or sneeze | 93.7 | 96.7 | 89.0 | 87.2 | 90.5 | 90.9 |
| Avoiding touching your eyes, nose, and mouth with unwashed hands | 90.9 | 92.7 | 90.1 | 88.0 | 91.0 | 90.5 |
| Use of disinfectants to clean hands when water and soap was not available for washing hands | 92.9 | 93.5 | 83.9 | 83.8 | 90.5 | 88.0 |
| Physical distancing | 84.1 | 85.4 | 85.9 | 79.5 | 83.1 | 84.3 |
| Disinfecting mobile phone | 84.2 | 82.1 | 83.4 | 84.6 | 83.6 | 83.6 |
| Disinfecting surfaces | 73.0 | 73.2 | 79.0 | 74.4 | 76.2 | 76.1 |
| Staying home when you were sick or when you had a cold | 63.1 | 65.9 | 66.9 | 61.5 | 62.4 | 64.6 |
| Total, n (%) | 252 (22.2) | 123 (10.8) | 453 (39.3) | 117 (10.3) | 189 (16.7) | 1134 (100) |

This study also investigated the attitude of the healthcare workers with regard to which group of people they recommend to use a facemask or N95 respirator. The vast majority of the respondents (94.8%) recommended the use of a facemask by all healthcare professionals, all healthy people to protect themselves from coronavirus infection (90.1%), and people with close contact with suspected or confirmed COVID-19 (88.8%). About 87% of all respondents suggested that N95 respirator should be used by all healthcare professionals as well as by people who are being in close contact with suspected or confirmed COVID-19 patients. About five in 10 (48%) of the respondents recommended the use of N95 respirator by healthy people to protect themselves against coronavirus infection. About 65% and 48% of the respondents from TASH and SPHMMC, respectively, recommended the use of N95 respirator for all healthy people to protect themselves from COVID-19.

### Exposure and preparedness in providing care to COVID-19 and other infectious disease outbreaks

Only about one-third (30.7%) of the study participated reported that they ever participated in direct clinical care to patients affected by infectious disease outbreaks such as Ebola, SARS and cholera. Nearly three in 10 (28.9%, n=328) respondents reported that they ever provided direct clinical care to at least one suspected/confirmed COVID-19 patient, with 39.1% participants from SPHMMC, 34.5% from MH and 31.1% from TASH. Regarding the level of preparedness of healthcare professionals to provide direct clinical care to COVID-19 patients, 33.6% (n=381) reported that they were prepared to provide direct clinical care to COVID-19 patients. In contrast, about two-third (66.4%) of the healthcare workers reported that they were not prepared to manage COVID-19 patients.

#### Risk perception of healthcare professionals due to their role in the COVID-19 pandemic

The study participants were asked questions about their personal health, potential risks of becoming infected with COVID-19 or the potential risks to their families and loved ones due to their clinical role in the hospital. About 30% and 43% of the participants somewhat or strongly worried, respectively, that their personal health is at risk during the COVID-19 pandemic due to their role in the hospital (Table 3). Nevertheless, 6% and 13.5% of respondents reported that they somewhat not worried or even not worried at all that their personal health was not at risk due to COVID-19. About 38% and 50% of all respondents perceived that they were somewhat worried or extremely worried about themselves, respectively, due to the potential risk of becoming infected with coronavirus by their clinical role in the hospital setting these days, with only 5.6% perceived that they were not worried about the risk of being infected with the virus. Majorities of the respondents (64.4%) extremely worried about the potential risk of infection to their family and loved ones, and the remaining 26.7% were somewhat worried. Only 4.4% of the respondents were not worried about the risk of COVID-19 to their family and loved ones.

**Table 3.** Healthcare professional’s worry about their clinical role in the hospital during COVID-19 by professional category (n=3 items)

| Variable | Professional category, % |  |  |  |  |
| --- | --- | --- | --- | --- | --- |
|  | *Physician*<br>(n=244) | *Intern*<br>(n=120) | *Nurse*<br>(n=431) | *Midwife*<br>(n=108) | *Other\**<br>(n=181) |
| How worried are you about your personal health due to your role in the hospital during COVID-19 pandemic? |  |  |  |  |  |
| Extremely worried | 47.1 | 50.0 | 39.7 | 40.7 | 42.0 |
| Somewhat worried | 35.2 | 27.5 | 25.5 | 28.7 | 37.0 |
| Average | 4.9 | 8.3 | 9.5 | 6.5 | 5.5 |
| Somewhat not worried | 3.7 | 5.0 | 7.9 | 7.4 | 4.4 |
| Not worried at all | 9.0 | 9.2 | 17.4 | 16.4 | 11.0 |
| How worried are you about the potential risk of becoming infected with COVID-19 due to your role in the hospital? |  |  |  |  |  |
| Extremely worried | 47.1 | 56.7 | 48.5 | 58.3 | 46.4 |
| Somewhat worried | 47.5 | 35.0 | 34.8 | 29.6 | 40.3 |
| Average | 3.3 | 6.7 | 8.6 | 5.6 | 6.6 |
| Somewhat not worried | 2.0 | 1.7 | 5.1 | 2.8 | 4.4 |
| Not worried at all | 0.0 | 0.0 | 3.0 | 3.7 | 2.2 |
| How worried are you about the potential risk COVID-19 to your family, loved ones or others due to your role in the hospital? |  |  |  |  |  |
| Extremely worried | 66.8 | 75.8 | 61.9 | 63.0 | 60.2 |
| Somewhat worried | 29.5 | 19.2 | 25.5 | 28.7 | 29.3 |
| Average | 2.5 | 4.2 | 7.4 | 4.6 | 5.0 |
| Somewhat not worried | 0.4 | 0.8 | 3.2 | 2.8 | 2.8 |
| Not worried at all | 0.8 | 0.0 | 1.9 | 0.9 | 2.8 |

The study participants were asked 12 questions to quantify their fears and worries (risk perception) about COVID-19 crisis, on a 3-point scale, where 1=don’t worry at all, 2=worry somehow and 3=worry a lot. Of the total 1134 study participants, 952 (84%) had complete responses on all the 12-items for computing the total score. About 66% of the respondents reported that they worried a lot about losing someone due to COVID-19, 66.7% worried a lot about the health of their loved ones, and 67.5% worried a lot about the health system being overloaded by the patients of COVID-19, followed by a lot of worries about the economic recession in the country (58%), and restricted access to food supplies (56.1%) (Table 4). The study also revealed that there were respondents who were ambivalent or didn’t worry at all about COVID-19 crisis.

**Table 4.** Healthcare professional’s fears and worries about COVID-19 crisis by hospital (n=12 items)

| Fear and worry question | Professional category, % |  |  |  |  | Total, %<br>(n=952) |
| --- | --- | --- | --- | --- | --- | --- |
|  | *Physician*<br>(n=221) | *Intern*<br>(n=110) | *Nurse*<br>(n=374) | *Midwife*<br>(n=95) | *Other\**<br>(n=152) |  |
| Loosing someone I love |  |  |  |  |  |  |
| Don't worry at all | 7.2 | 10.0 | 12.6 | 15.8 | 12.5 | 11.3 |
| Worry somehow | 25.3 | 19.1 | 21.1 | 22.1 | 23.7 | 22.4 |
| Worry a lot | 67.4 | 70.4 | 66.3 | 62.1 | 63.8 | 66.3 |
| Health system being overloaded |  |  |  |  |  |  |
| Don't worry at all | 7.7 | 5.5 | 8.8 | 9.5 | 9.2 | 8.3 |
| Worry somehow | 17.2 | 30.0 | 24.1 | 29.5 | 27.0 | 24.2 |
| Worry a lot | 75.1 | 64.5 | 67.1 | 61.1 | 63.8 | 67.5 |
| My own mental health |  |  |  |  |  |  |
| Don't worry at all | 19.9 | 30.0 | 22.2 | 21.1 | 25.0 | 22.9 |
| Worry somehow | 44.8 | 30.9 | 35.6 | 37.9 | 36.2 | 37.5 |
| Worry a lot | 35.3 | 39.1 | 42.2 | 41.1 | 38.8 | 39.6 |
| My own physical health |  |  |  |  |  |  |
| Don't worry at all | 11.8 | 12.7 | 17.4 | 11.6 | 17.8 | 15.0 |
| Worry somehow | 45.2 | 40.9 | 38.8 | 45.3 | 38.2 | 41.1 |
| Worry a lot | 43.0 | 46.4 | 43.9 | 43.2 | 44.1 | 43.9 |
| My loved ones' health |  |  |  |  |  |  |
| Don't worry at all | 12.7 | 8.2 | 10.2 | 10.5 | 9.2 | 10.4 |
| Worry somehow | 21.3 | 14.5 | 26.5 | 26.3 | 20.4 | 22.9 |
| Worry a lot | 66.1 | 77.3 | 63.4 | 63.2 | 70.4 | 66.7 |
| Restricted liberty of movement |  |  |  |  |  |  |
| Don't worry at all | 13.6 | 18.2 | 12.8 | 13.7 | 13.2 | 13.8 |
| Worry somehow | 44.8 | 41.8 | 43.9 | 43.2 | 49.3 | 44.6 |
| Worry a lot | 41.6 | 40.0 | 43.3 | 43.2 | 37.5 | 41.6 |
| Small companies running out of business |  |  |  |  |  |  |
| Don't worry at all | 10.9 | 13.6 | 14.2 | 16.8 | 12.5 | 13.3 |
| Worry somehow | 50.2 | 50.0 | 37.2 | 35.8 | 38.2 | 41.7 |
| Worry a lot | 38.9 | 36.4 | 48.7 | 47.4 | 49.3 | 45.0 |
| Economic recession in my country |  |  |  |  |  |  |
| Don't worry at all | 7.7 | 7.3 | 9.1 | 7.4 | 10.5 | 8.6 |
| Worry somehow | 37.1 | 47.3 | 29.7 | 36.8 | 25.0 | 33.4 |
| Worry a lot | 55.2 | 45.5 | 61.2 | 55.8 | 64.5 | 58.0 |
| Restricted access to food supplies |  |  |  |  |  |  |
| Don't worry at all | 11.3 | 5.5 | 9.9 | 6.3 | 8.6 | 9.1 |
| Worry somehow | 37.1 | 35.5 | 31.6 | 36.8 | 37.5 | 34.8 |
| Worry a lot | 51.6 | 59.1 | 58.6 | 56.8 | 53.9 | 56.1 |
| Becoming unemployed |  |  |  |  |  |  |
| Don't worry at all | 51.1 | 27.3 | 25.4 | 22.1 | 28.9 | 31.8 |
| Worry somehow | 19.5 | 27.3 | 32.9 | 32.6 | 26.3 | 28.0 |
| Worry a lot | 29.4 | 45.5 | 41.7 | 45.3 | 44.7 | 40.1 |
| Not being able to pay my bills |  |  |  |  |  |  |
| Don't worry at all | 30.8 | 23.6 | 18.2 | 17.9 | 17.1 | 21.5 |
| Worry somehow | 37.1 | 33.6 | 42.2 | 43.2 | 41.4 | 40.0 |
| Worry a lot | 32.1 | 42.7 | 39.6 | 38.9 | 41.4 | 38.4 |
| Unable to visit people who depend on me |  |  |  |  |  |  |
| Don't worry at all | 10.9 | 14.5 | 8.8 | 4.2 | 12.5 | 10.1 |
| Worry somehow | 32.6 | 29.1 | 34.8 | 40.0 | 27.6 | 33.0 |
| Worry a lot | 56.6 | 56.4 | 56.4 | 55.8 | 59.9 | 56.9 |

An overall fear and worry index about COVID-19 was created using 12 questions. The overall score for the scale was calculated by summing up the score of all questions (from 12 to 36). The higher the score, the greater the fear and worry of the COVID-19. Table 5 presents the mean scores for each and the overall worry indicators of COVID-19 crisis by professional category. Overall, the participants reported an average of moderate-to-high levels of COVID-19 worry (2.37) on each item, ranging from 2.1 on ‘becoming unemployed’ to 2.6 on ‘losing someone they love’, ‘health system being overloaded’ and ‘someone’s loved health’. The overall average worry score of the 12 items for the COVID-19 crisis was high, with a mean (±SD) of 28.4 (±5.9), ranging from 12 to 36. The total average fear and worry scores for the hospitals ranged from 25.6 (±6.8) at TASH to 31.3 (±5.0) at GMH; and was further categorized into three levels i.e. low, moderate, and high fear and worry level. Figure 1 shows the pattern of the total fear and worry scores of COVID-19 crisis, and about 56% of respondents from TASH showed a relatively low fear and worry score compared to the highest (50.9%) fear and worry score reported by participants from GMH.

**Table 5.** Mean fear and worry scores of healthcare professionals about COVID-19 crisis by professional category (n=12 items)

| COVID-19 worry items | Professional category, Mean (SD)* |  |  |  |  | Mean (SD)<br>(n=952) |
| --- | --- | --- | --- | --- | --- | --- |
|  | *Physician*<br>(n=221) | *Intern*<br>(n=110) | *Nurse*<br>(n=374) | *Midwife*<br>(n=95) | *Other\**<br>(n=152) |  |
| Losing someone I love | 2.6 (0.6) | 2.6 (0.7) | 2.5 (0.7) | 2.5 (0.8) | 2.5 (0.7) | 2.6 (0.7) |
| Health system being overloaded | 2.7 (0.6) | 2.6 (0.6) | 2.6 (0.6) | 2.5 (0.7) | 2.6 (0.7) | 2.6 (0.6) |
| My own mental health | 2.2 (0.7) | 2.1 (0.8) | 2.2 (0.8) | 2.2 (0.8) | 2.1 (0.8) | 2.2 (0.8) |
| My own physical health | 2.3 (0.7) | 2.3 (0.7) | 2.3 (0.7) | 2.3 (0.7) | 2.3 (0.7) | 2.3 (0.7) |
| My loved one's health | 2.5 (0.7) | 2.7 (0.6) | 2.5 (0.7) | 2.5 (0.7) | 2.6 (0.7) | 2.6 (0.7) |
| Restricted liberty of movement | 2.7 (0.7) | 2.2 (0.7) | 2.3 (0.7) | 2.3 (0.7) | 2.2 (0.7) | 2.3 (0.7) |
| Companies running out of business | 2.3 (0.7) | 2.2 (0.7) | 2.4 (0.7) | 2.3 (0.7) | 2.4 (0.7) | 2.3 (0.7) |
| Economic recession in my country | 2.5 (0.6) | 2.4 (0.6) | 2.5 (0.7) | 2.5 (0.6) | 2.5 (0.7) | 2.5 (0.7) |
| Restricted access to food supplies | 2.4 (0.7) | 2.5 (0.6) | 2.5 (0.7) | 2.5 (0.6) | 2.6 (0.6) | 2.5 (0.7) |
| Becoming unemployed | 1.8 (0.9) | 2.2 (0.8) | 2.2 (0.8) | 2.2 (0.8) | 2.2 (0.8) | 2.1 (0.8) |
| Not being able to pay my bills | 2.0 (0.8) | 2.2 (0.8) | 2.2 (0.7) | 2.2 (0.7) | 2.2 (0.7) | 2.2 (0.8) |
| Not able to visit people | 2.5 (0.6) | 2.4 (0.7) | 2.5 (0.7) | 2.5 (0.6) | 2.5 (0.7) | 2.5 (0.7) |
| Overall mean (SD) | 27.9 (5.9) | 28.5 (5.6) | 28.7 (6.1) | 28.6 (5.8) | 28.6 (5.7) | 28.4 (5.9) |

\*Numbers in parentheses represent standard deviations.

**Fig. 1.**
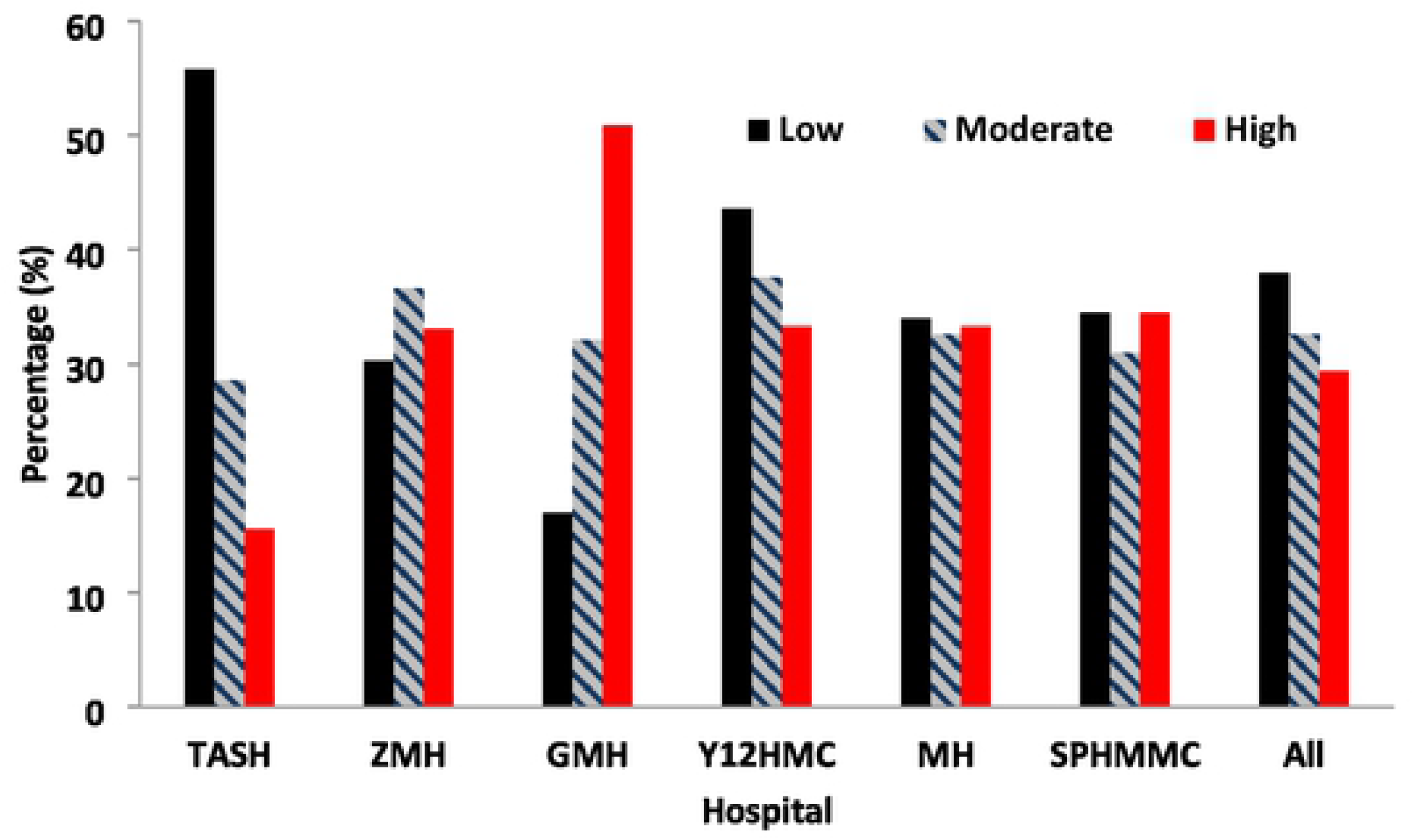
Pattern of fear and worry scores of COVID-19 crises by hospital

The total fear and worry scores of COVID-19 was finally changed into binary using the Visual Binning in SPSS (low fear/worry ≤29 and high fear/worry >29 score). Table 6 shows the results of bivariate and multivariable logistic regression analyses of predictors associated with respondents mean scores of fears and worries about COVID-19 crisis. In the bivariate analyses departments/units and the hospitals were significantly associated with fear and worry scores of COVID-19 crises. Nurses were 1.52 times more likely to report fear and worry (OR=1.52, 95% CI:1.09-2.13, *P*<0.015), and healthcare workers who ever participated in clinical care to Ebola, SARS and cholera patients were 0.67 times less likely to report fear and worry due to COVID-19 crisis (OR=1.67, 95% CI:0.51-0.88, *P*<0.005).

**Table 6.** Factors associated with worries about COVID-19 crisis in the study population using multiple logistic regression analyses (n=952)

| Predictor | Fear and worry level, n (%) |  | Crude OR (95% CI)* | P-value | Adjusted OR (95% CI) | P-value |
| --- | --- | --- | --- | --- | --- | --- |
| | Low ($\leq 29$) | High ($> 29$) | | | | |
| Gender |  |  |  |  |  |  |
| Male | 255 (55.8) | 202 (44.7) | 0.84 (0.65-1.09) | 0.186 | 0.96 (0.73-1.28) | 0.792 |
| Female | 255 (51.5) | 240 (48.5) | 1.0 |  | 1.0 |  |
| Professional category |  |  |  |  |  |  |
| Physician | 131 (59.3) | 90 (40.7) | 1.0 |  | 1.0 |  |
| Intern | 61 (55.4) | 49 (44.5) | 1.17 (0.74-1.86) | 0.507 | 0.78 (0.47-1.30) | 0.336 |
| Nurse | 183 (48.9) | 191 (51.1) | 1.52 (1.09-2.13) | 0.015 | 1.33 (0.91-1.93) | 0.139 |
| Midwife | 54 (56.8) | 41 (43.2) | 1.11 (0.68-1.80) | 0.687 | 0.69 (0.37-1.26) | 0.226 |
| Other*** | 81 (53.3) | 71 (46.7) | 1.28 (0.84-1.94) | 0.252 | 1.37 (0.83-2.24) | 0.218 |
| Department/Unit |  |  |  |  |  |  |
| Gyn&Ob | 78 (45.9) | 92 (54.1) | 1.0 |  |  |  |
| Surgical | 75 (59.1) | 52 (40.9) | 0.59 (0.37-0.94) | 0.025 | 0.65 (0.37-1.16) | 0.142 |
| Pediatrics | 69 (51.9) | 64 (48.1) | 0.79 (0.50-1.24) | 0.300 | 0.82 (0.47-1.44) | 0.492 |
| Medical | 74 (60.2) | 49 (39.8) | 0.56 (0.35-0.90) | 0.016 | 0.63 (0.35-1.13) | 0.119 |
| OPD/Screening/Triage | 57 (47.9) | 62 (52.1) | 0.92 (0.58-1.48) | 0.735 | 0.84 (0.47-1.50) | 0.546 |
| Emergency | 55 (68.8) | 25 (31.3) | 0.39 (0.22-0.68) | 0.001 | 0.40 (0.21-0.77) | 0.006 |
| Anesthesia/OR/IC | 31(40.8) | 45 (59.2) | 1.23 (0.71-2.13) | 0.458 | 1.11 (0.57-2.16) | 0.761 |
| Other*** | 71 (57.3) | 53 (42.7) | 0.63 (0.40-1.01) | 0.055 | 0.52 (0.28-0.96) | 0.0.8 |
| Hospital |  |  |  |  |  |  |
| TASH | 154 (68.8) | 70 (31.1) | 0.44 (0.30-0.66) | <0.001 | 0.49 (0.32-0.75) | 0.001 |
| ZMH | 64 (45.1) | 78 (54.9) | 1.18 (0.77-1.82) | 0.448 | 1.34 (0.85-2.11) | 0.209 |
| GMH | 38 (33.9) | 74 (66.1) | 1.89 (1.17-3.06) | 0.009 | 1.87 (1.10-3.18) | 0.020 |
| Y12HMC | 88 (62.2) | 45 (33.8) | 0.50 (0.34-0.78) | 0.003 | 0.52 (0.32-0.84) | 0.008 |
| MH | 69 (47.9) | 75 (52.1) | 1.05 (0.69-1.62) | 0.809 | 1.12 (0.72-1.75) | 0.607 |
| SPHMMC | 97 (49.2) | 100 (50.8) | 1 |  | 1 |  |
| Prepared to provide direct care to COVID-19 cases |  |  |  |  |  |  |
| Yes | 165 (49.5) | 168 (50.5) | 1.28 (0.98-1.67) | 0.068 | 1.04 (0.78-1.40) | 0.776 |
| No | 345 (55.7) | 274 (44.3) | 1.0 |  | 1.0 |  |
| Ever provided clinical care to suspected/confirmed COVID-19 patients |  |  |  |  |  |  |
| Yes | 147 (50.3) | 145 (49.7) | 1.21 (0.92-1.59) | 0.184 | 1.34 (1.02-1.91) | 0.037 |
| No | 363 (55.0) | 297 (45.0) | 1.0 |  | 1.0 |  |
| Ever participated in clinical care to Ebola, SARS and cholera patients |  |  |  |  |  |  |
| Yes | 181 (60.3) | 119 (39.7) | 0.67 (0.51-0.88) | 0.005 | 0.66 (0.48-0.90) | 0.009 |
| No | 329 (50.5) | 323 (49.5) | 1.0 |  | 1.0 |  |

In the multivariable logistic regression analyses, hospitals retained the statistical significance for the fear and worry score, where respondents from TASH (adjusted OR=0.49, 95% CI:0.32-0.75, *P*=0.001) and Y12HMC (adjusted OR=0.52, 95% CI:0.32-0.84, *P*=0.008) were less likely to report fear and worry about COVID-19 crisis (Table 6). In contrast, respondents from GMH were more likely to fear and worry for COVID-19 crisis (adjusted OR=1.77, 95% CI:1.10-3.18) than those from the SPHMMC respondents. Healthcare professionals ever provided clinical care to suspected/ confirmed COVID-19 patients were 1.34 times more likely to report fear and worry due to COVID-19 crises (OR=1.34, 95% CI:1.02-1.91, *P*=0.037), however respondents who ever participated in clinical care to Ebola, SARS and cholera patients were 0.66 times less likely to report fear and worry due to COVID-19 crisis (OR=0.66, 95% CI:0.48-0.90, *P*=0.009). Gender, professional category and preparedness to provide direct care to COVID-19 patients did not appear significant in the multivariable logistic regression model to predict the odds of fear and worry score for COVID-19 crisis.

## Discussion

Since its emergence in December 2020, the COVID-19 pandemic is a global public health concern and the most current topic of discussion across every facet of life, especially among the healthcare professionals and patients. This study was conducted in Addis Ababa city during 09-26 June 2020, three months after detection of the first confirmed case of COVID-19 in Ethiopia. Addis Ababa city is the most affected part in the country. The study aimed to assess the risk perceptions and protective behaviors of COVID-19 among healthcare professionals in the city. Our study participants include medical doctors, interns, nurses, midwives, pharmacists, medical laboratory technologists, and technicians. These categories of healthcare professionals have direct or indirect close personal exposures with suspected or confirmed COVID-19 patients while performing their clinical duties.

The overwhelming majority of the participants in our study reported a high level of practice towards the prevention of COVID-19 infection particularly regarding using facemask, hand washing for at least 20 seconds, covering mouth and nose when coughing or sneezing, and avoiding touching eyes, nose, and mouth with unwashed hands as far as possible. This finding is consistent with the finding of a similar study conducted in China, where the risk of spread of COVID-19 has largely improved the infection prevention and control behaviors of healthcare professionals working in hospitals [18]. In a study conducted in Egypt, hand washing, refraining from touching eyes, mouth and nose, and using surgical facemask were the most frequently accepted preventive measures among health workers [19]. The WHO recommends the use of primary preventive measures that includes regular hand washing, social distancing, and respiratory hygiene (covering mouth and nose while coughing or sneezing) by healthcare workers in order to prevent the spread of the virus among themselves and patient’s close contacts [20].

Studies conducted during the early stage of the pandemic revealed that healthcare workers had insufficient knowledge about COVID-19 pandemic to protect themselves from coronavirus infection [21]. In one study in Greece, only 25% of healthcare practitioners washed their hands after touching a patient, despite the fact that 94% of the respondents knew that SARS-CoV-2 transmission could be reduced with hand washing [22]. Although hand washing is recommended for the general public in order to prevent the transmission of COVID-19, hand hygiene is mandatory for health care practitioners, in order to prevent infections, both for oneself and for the patients [23]. In the present study, the use of facemask was reported to be 93%. A recent study conducted in Addis Ababa just before our study revealed that about two-third of the healthcare workers demonstrated a poor practice of facemask utilization [24]. Similar results were reported in North-East India that majority of the healthcare workers (91%) reported that they used surgical masks, 97% were using hand sanitizer and 97% participants were properly using hand hygiene [25].

In the present study, the majority of the study participants recommended mask-wearing by all healthcare professionals, all healthy people to protect themselves from coronavirus infection, and people with close contact with suspected or confirmed COVID-19. Similarly, about 87% of the respondents suggested that N95 respirator should be used by all healthcare professionals as well as by people who are being in close contact with suspected or confirmed COVID-19 patients. In Pakistan, 71% of the healthcare workers believed that wearing general medical masks was protective against COVID-19 [26], and studies also suggested that surgical masks are similarly as effective as N95 respirators if used with hand wash and other infection prevention precautions [27]. However, a rapid systematic review on the efficacy of facemasks and respirators against coronaviruses and other respiratory transmissible viruses reported that continuous use of respirators is more protective compared to the medical masks, and medical masks are more protective than cloth masks among health workers in healthcare settings [28].

This study demonstrated that about one-third of all respondents in our study either participated in direct clinical care to patients affected by an infectious disease outbreak (e.g., Ebola virus, SARS, cholera, Zika virus) (31%) or provided direct clinical care at least for one suspected or confirmed COVID-19 patients (29%) during the current COVID-19 epidemic. This percentage is higher from other studies on this subject in the early days of the COVID-19 outbreak in China [29]. A significant number (38%) of healthcare professionals in the current study expressed lack of or low level of preparedness to manage suspected or confirmed COVID-19 patients. This raises a concern regarding the ability and confidence of the healthcare workers to combat COVID-19 infection. Despite these concerns, along with the shortage of PPE and inadequate training during the COVID-19, the healthcare workers continue to work with the management of suspected or confirmed COVID-19, working in the hospital setting where COVID-19 patients were admitted, risking their lives to save their patients. However, this could highlight the risk of infection among healthcare workers and cross-contamination within hospitals and could lead to a higher rate of hospital-acquired infections. Therefore, our study provides considerable insights into the necessity of immediate and determined efforts focused on training programs and providing an adequate supply of PPE to ensure the safety of health personnel during the COVID-19 pandemic [30].

In the present study, about 88% of the healthcare professionals were afraid of being infected with the disease and about 91% were worried about the potential risk of transmitting the virus to their family and loved ones. The risk of contracting the virus was perceived to be very high at the time of the study. Healthcare workers expressed worry and fear of infection due to the contagious nature of the virus, close contact with suspected and confirmed COVID-19 patients, and infection happening to their family and colleagues. In Iran, it was found that about 92% of the healthcare workers worried about being infected with the virus and transmitting it to the family [31]. In a study conducted in Henan province of China, 89% of healthcare workers had sufficient knowledge of COVID-19, 85% were concerned about infection with the virus, and 90% followed correct practices regarding the prevention of COVID-19 [32]. About 83% of the healthcare workers in Egypt reported increased risk perception because of the concern of being infected with COVID-19 and fear of transmitting the disease to their families, and 89% stated that they were more susceptible to COVID-19 infection mainly due to the shortage of PPE [19].

In the current study, the overall risk perception expressed in fear and worry score of the study participants regarding COVID-19 crisis was considerably higher, with a mean of 28, ranging from 12 to 36. Various studies have reported the psychological impact of COVID-19 on healthcare professionals [33]. A recent scoping review found that the frontline healthcare workers are at an increased risk of direct physical and mental consequences as the result of providing care to patients with COVID-19 [34]. Studies demonstrated that more than 50% of healthcare professionals report symptoms of depression, insomnia, and anxiety due to COVID-19 [35]. A recent study carried out in Pakistan on fear and anxiety among healthcare professionals reported that about three-fourth of them had fear of getting infected during the management of COVID-19 patients, and another two-third reported severe anxiety, which was particularly more common among nurses [36]. Studies also reported excessive workload, isolation, mental stress and discrimination among frontline health professionals, thus, contributing to physical exhaustion, emotional disturbance, worry and fear [37]. A Cochrane review reported the suffering of healthcare workers from work-related or occupational stress, which can be reduced by cognitive-behavioral training as well as mental and physical relaxation [38]. A multicenter study conducted among frontline nurses in China showed poor mental health during the COVID-19 outbreak, mainly due to the fear of contracting the virus and high workload [39]. Moreover, the same study revealed that nurses who were confident in their infection control knowledge and skills had lower stress levels than those who felt less prepared.

Finally, this study had several limitations. First, the study had a potential to be affected by selection bias and eligible participants might be excluded. Second, this study was conducted in six public hospitals in Addis Ababa, and may possibly limit the generalization of the results and findings to other public and private hospitals. Third, the study focused on more general populations of healthcare professionals similar to other studies [32,40] rather than healthcare workers who might have direct contact with COVID-19 patients [41]. Finally, the results of this study are based on self-reported data, and the respondents may overestimate or underestimate the responses in a way that they believe is socially acceptable rather than reporting actual or genuine answers. Despite these limitations, the results obtained provide important information to guide health communication efforts that can support prevention efforts of COVID-19 among healthcare professionals.

## Conclusions

In conclusion, our study has illuminated the current level of risk perception and preventive practices of COVID-19 among healthcare professionals, with a special focus on those working in the clinical departments of the hospitals who have direct or indirect contact with COVID-19 patients. The present study findings demonstrated that healthcare professionals participated in the study showed a universally higher preventive practices to prevent COVID-19 infections. The healthcare workers perceived high level of COVID-19 risk particularly due to shortage of PPE, and majority reported that they didn’t receive any training in infection prevention and control measures since COVID-19, although they had adequate level of practice to protect themselves from the infection of novel coronavirus. Likewise, majority of the participants reported that they worried about the potential risk of becoming infected with COVID-19 and transmitting the disease to their family. The present study also was able to identify factors associated with fear and worry related to COVID-19 crisis in order to address them during the implementation of risk communication programs with the public and healthcare during the current COVID-19 pandemic.

## Acknowledgments

The authors are grateful to the participating hospitals and their healthcare staff for committing their time and voluntarily filling in the questionnaire. They are also thankful to all data collectors and logistics facilitators for their time and commitment.

## Ethical approval and consent to participate

Ethical clearance was obtained from the Institutional Review Board of the College of Health Sciences at Addis Ababa University. All participants gave their informed consent.

## Contributors

Conceptualization: WD, AW, WAA, WA

Designing of the study: WD, AW, MG, WAA, WA

Data curation: WD, AW

Statistical analysis: WD, AW

Field supervision: MG, WD, AW

Writing – original draft: WD, AW

Writing – review & editing: WD, AW, MG, WAA, WA

Substantial contribution to the interpretation of the data: WD, AW, MG, WAA, WA

All the authors read and approved the final manuscript.

